# Ryder: Epigenome normalization using a two-tier model and internal reference regions

**DOI:** 10.64898/2026.03.15.711886

**Authors:** Yaqiang Cao, Guangzhe Ge, Keji Zhao

**Author notes:** These authors contributed equally to this work (Yaqiang Cao, Guangzhe Ge).

## Abstract

**Motivation:** Sequencing-based epigenomic profiling methods are powerful but suffer from technical variability that complicates cross-sample comparisons and can obscure true biological signals. While existing normalization methods using spike-in controls or computational approaches have been proposed, they often rely on assumptions that may not hold across diverse experimental conditions or require additional data types.

**Results:** We present Ryder, a flexible and robust Python package for the normalization of epigenomic signal tracks. Ryder leverages stable internal reference regions, such as invariant CTCF binding sites, to correct for technical artifacts genome-wide. Our results show that it effectively adjusts both background noise and signal intensity, ensuring accurate signal alignment across samples while preserving genuine biological differences. We demonstrate that Ryder performs robustly across diverse assays including DNase-seq, CUT&RUN, ATAC-seq, MNase-seq, and ChIP-seq, with or without spike-in controls. By reducing technical noise, Ryder improves the detection of genuine biological changes, such as quantitative reduction of chromatin accessibility at key enhancer elements by depletion of BRG1, a key subunit of the chromatin remodeling BAF complexes.

**Availability and Implementation:** The Ryder source code, documentation and test data are freely available at: https://github.com/YaqiangCao/ryder. The software version used in this study is archived at Zenodo: https://zenodo.org/records/21267457.

## 1 Introduction

Sequencing-based epigenomic methods such as ChIP-seq (Barski, et al., 2007; Park, 2009), DNase-seq (He, et al., 2014; Song and Crawford, 2010), ATAC-seq (Buenrostro, et al., 2013; Grandi, et al., 2022), and MNase-seq (Henikoff, et al., 2011; Schones, et al., 2008) have revolutionized our understanding of chromatin structure and gene regulation. Large-scale projects have generated comprehensive maps of DNA-protein interactions, chromatin accessibility, and nucleosome positioning, highlighting the complexity of gene regulatory networks (Consortium, 2012; Kundaje, et al., 2015; Martens and Stunnenberg, 2013; Stunnenberg, et al., 2016), further advocating the application of these methods in addressing fundamental biological questions. However, technical variability—stemming from sample quality, library preparation, sequencing depth, and batch effects (Meyer and Liu, 2014) — remains a significant obstacle, often manifesting as inconsistent signal-to-noise ratios between samples. Without appropriate normalization, these artifacts can obscure biological insights, resulting in misinterpretation.

Both experimental and computational normalization methods aimed at mitigating technical biases have been proposed. Spike-in controls, which incorporate exogenous chromatin or cells, provide a direct reference but depend critically on the assumption of equivalent experimental conditions between spike-in and endogenous materials — an assumption that is difficult to validate and may not hold across diverse conditions or protocols. For instance, a recent study found that even minor, unintentional variations in the amount of spike-in DNA added to each sample led to significant, artifactual differences in read counts, making the spike-in data unusable for normalization (Hagihara, et al., 2025). Additionally, optimal titration of the spike-in material is critical: too little limits its utility, while too much can consume sequencing capacity (Patel, et al., 2024). Furthermore, a single scaling factor derived from spike-ins provides a global normalization, but it may not fully capture localized or site-specific variations (Egan, et al., 2016). On the computational side, methods such as MAnorm operate on the assumption that shared peaks between ChIP-seq samples reflect regions of invariant bindings and should therefore exhibit comparable signal intensities (Shao, et al., 2012). However, this assumption becomes questionable when there are quantitative differences within these shared peaks, and it may not hold true when global changes are present, such as during cellular aging (Bozukova, et al., 2022; Cheng, et al., 2018; Morandini, et al., 2024). More recent methods, such as S3norm, attempt to account for both sequencing depth and signal-to-noise ratio through iterative modeling (Xiang, et al., 2020). However, its effectiveness can depend on accurate segmentation of the genome into distinct signal and background regions. This clear separation may be challenging for broadly spread signals, such as those often observed in H3K27me3 or H3K9me3 ChIP-seq data. Other strategies, such as IGN (Hu, et al., 2025), rely on a set of presumed invariantly expressed genes to anchor normalization. However, identifying such stable loci requires ideally matched RNA-seq data, which is assumed to be free of technical variation—an assumption that may not always hold.

Given these challenges, a flexible normalization framework for epigenomic data, capable of accommodating diverse experimental designs, technical variability, and biological contexts, and allowing for easily testable assumptions, is essential. We developed Ryder, a flexible tool designed for cross-sample normalization and the identification of variable features. Ryder leverages stable internal reference regions—such as invariant CTCF sites or user-defined transcription start sites (TSSs) where a reasonable assumption of stability can be tested—to normalize both background and signal genome-wide, aiming for consistent signal alignment and signal-to-noise ratios. Its default two-tier strategy separates background correction from signal alignment, and it also supports single-scaling or parameter estimation from spike-in controls. This flexible design enables Ryder to normalize a range of assays—including DNase-seq, CUT&RUN, ATAC-seq, MNase-seq, and ChIP-seq—with or without spike-ins, making it potentially effective for both subtle perturbations and global chromatin shifts.

## 2 Methods

### 2.1 Overview of Ryder

Within the Ryder package, normalization and differential feature detection are performed sequentially by paw.py and patrol.py, with bigWig files as primary inputs. Paw.py rescales the target bigWig signals using stable internal reference regions, corrects background noise via distribution-based modeling, and outputs both quality-control metrics and normalized signal tracks. Patrol.py then analyzes these tracks across user-defined regions, applying fold-change or Poisson-based statistics to identify differential features, generate M (log ratio) A (average) plots, and export BED files for downstream interpretation.

The primary input to paw.py is a pair of bigWig files representing a reference/control sample and a target/treatment sample. These bigWig files are expected to be pre-normalized by total sequencing depth, such as reads per million mapped reads (RPM) or counts per million mapped reads (CPM). Ryder then estimates additional normalization parameters to correct differences in background signal, foreground/reference-region signal, and signal-distribution shape between the two samples. The main output of paw.py is a normalized target bigWig file aligned to the reference sample, together with diagnostic plots and estimated normalization parameters.

### 2.2 Internal reference regions

Ryder accepts any user-provided BED file of putatively stable genomic regions. We assume that constitutive CTCF binding sites—conserved across samples and overlapping with peaks in the dataset—can serve as internal reference regions. This is supported by CTCF’s well-established role as a ubiquitous and stable chromatin architectural protein across diverse cell types and conditions (Dixon, et al., 2012; Fang, et al., 2020; McArthur and Capra, 2021). Importantly, this assumption is also readily testable, particularly in cases where the perturbed factor is unrelated to CTCF and does not share its DNA-binding motif. Ryder provides human constitutive CTCF binding sites curated from a previous study (Fang, et al., 2020), as well as mouse invariant CTCF sites derived from CTCF ChIP-seq peaks curated in the Cistrome database (L’Yi, et al., 2023), defined as being consistently shared across more than 50 distinct cell and tissue types. To minimize potential artifacts from problematic genomic loci, regions overlapping ENCODE blacklist regions (Amemiya, et al., 2019) were removed from the preprocessed reference-region sets. These filtered reference files are provided in the Ryder GitHub repository: https://github.com/YaqiangCao/ryder/tree/main/refData.

For each analysis, the invariant reference set was intersected with assay-specific signal regions, such as DNA hypersensitive sites (DHSs), ATAC-seq peaks, CUT&RUN peaks, or ChIP-seq peaks, to ensure that normalization parameters were estimated from reference-associated regions with detectable signal in the corresponding assay. Unless otherwise specified, we retained the coordinates of the assay-specific signal regions that overlapped invariant reference sites using BEDtools (Quinlan and Hall, 2010): intersectBed -a targetRegion.bed -b reference.bed > ref.bed. Thus, the resulting ref.bed contains assay-specific intervals, for example DHSs or peaks, selected by overlap with the stable reference set. When the biological perturbation may affect a subset of candidate reference sites, those sites should be excluded before normalization.

Ryder is not restricted to CTCF sites. Users may provide alternative stable reference sets, such as transcription start sites of stably expressed housekeeping genes or binding sites of other factors, provided that the assumption is appropriate for the biological system and can be evaluated from the data. CTCF sites should not be used as the default reference set when the perturbation directly affects CTCF. In such cases, users should select an alternative stable reference set or provide spike-in-derived normalization parameters when suitable spike-in data are available.

### 2.3 Normalization framework

Let *c*_*i*_ and *t*_*i*_ denote the RPM normalized bigWig signal values for the control and treatment samples, respectively, at genomic interval *i* with the assumed same signal value. Ryder estimates five parameters for normalization: a background scaling factor *s*_*b*_, a foreground/reference-region scaling factor *s*_*f*_, two log_2_-space signal-alignment parameters *α* and *β*, and a linear-scale cutoff *τ*. In the default two-tier mode, the target signal is transformed as:

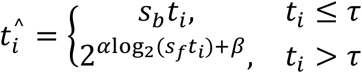

where 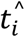 is the Ryder-normalized target signal. This two-tier strategy allows Ryder to correct background and foreground signal separately, rather than applying a single genome-wide scaling factor to all intervals.

By default, Ryder estimates all five parameters *s*_*b*_, *s*_*f*_, *α, β* and *τ* directly from the input data. Users can also bypass parameter estimation through the -pred option to specify *s*_*b*_, *s*_*f*_, *α, β* and -noise for *τ*. This flexibility is useful when normalization parameters are estimated from spike-in data and then applied to endogenous genomic tracks.

When spike-in controls were available, we also compared two spike-in-based normalization strategies: spike-in read-ratio normalization and spike-in profile normalization. For spike-in read-ratio normalization, the fraction of reads mapped to the spike-in genome was calculated for each condition as:

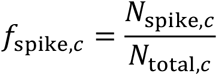

where *N*_spike,*c*_ is the number of mapped reads from condition *c* mapped to the spike-in genome, and *N*_total,*c*_ is the corresponding total library read count. The target scaling factor was then calculated relative to the reference condition:

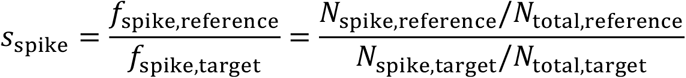

The target signal was normalized as: 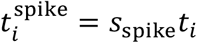. This single-factor normalization was done by running paw.py with -pred *s*_spike_ *s*_spike_ 1 0.

For spike-in profile normalization, the RPM-normalized signal tracks from the spike-in genome were analyzed with paw.py first using spike-in DHSs or peaks as reference regions to estimate four parameters: the background scaling factor 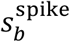, the foreground/reference-region scaling factor 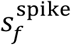, and the signal-distribution alignment parameters *α*^spike^ and *β*^spike^. These parameters were then transferred to the endogenous normalization by running paw.py with -pred 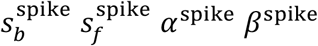.

Thus, spike-in read-ratio normalization applies one global factor derived from the relative abundance of spike-in reads, whereas spike-in profile normalization transfers separate background, foreground, and signal-distribution parameters estimated from the spike-in signal profiles.

### 2.4 Outlier removal from internal reference regions

Because not all candidate reference regions are guaranteed to remain stable in a given experiment, Ryder first removes outlier reference sites. For each candidate reference region *r*, Ryder computes the average bigWig signal in the reference and target samples. Ryder constructs an M-A representation:

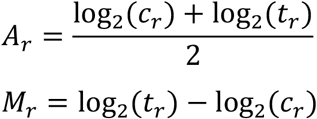

where *c*_*r*_and *t*_*r*_are the average signals over reference region *r*. Then Ryder computes the Mahalanobis distance (De Maesschalck, et al., 2000) of each region in the two-dimensional (*A*_*r*_, *M*_*r*_) space:

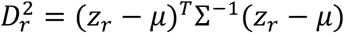

where *z*_*r*_ = (*A*_*r*_, *M*_*r*_), *μ* is the mean vector, and Σ is the covariance matrix estimated from all candidate reference regions. A Chi-square-based *p*-value is assigned to each region, and regions with *P* ≤ *p*_cut_ are removed as outliers. The default cutoff in paw.py is *p*_cut_ = 0.1. The remaining regions are used for estimating signal-alignment parameters *α* and *β*.

The Mahalanobis-derived p-value is used only to screen candidate reference regions before parameter estimation; it is not used to declare biological discoveries. Therefore, this p-value cutoff is a quality-control tuning parameter rather than a false-discovery-rate threshold, and larger p-value cutoff values remove more candidate regions. Users may decrease this p-value cutoff (-pcut option in paw.py) to retain more regions and should verify that sufficient reference regions remain and that the estimated parameters are stable.

### 2.5 Definition of foreground and background regions

After outlier removal, Ryder defines foreground regions as the retained reference regions. Background regions are generated by shifting each foreground region upstream and downstream by multiples of its own width. In the current implementation of paw.py, shifts of 1, 2, 3, 4, 5, and 10 region widths are used. The default shift multipliers were chosen to sample local flanking background at multiple distances: smaller shifts capture nearby local background, whereas the 10-width shift provides a more distal background estimate. Candidate background regions are discarded if they overlap the foreground regions or fall outside the coordinate range covered by reference regions on the same chromosome. These foreground and background regions are then used to estimate signal level, background level, and signal-to-noise differences between the control and treatment samples. Because broad or diffuse signals may extend into the flanks, these defaults should not be treated as universally valid. Users should inspect the noise-cutoff plot; when the two components are not adequately separated, an alternative reference set or an appropriate single-tier mode (-flat fg or -flat bg) should be used.

### 2.6 Estimation of scaling factors

Ryder computes binned average bigWig signals over foreground and background regions for both samples, using 100 bins per region by default. Let 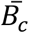 and 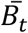 (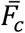 and 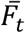) denote the average background (foreground) signals for the control and target samples, respectively. Ryder estimates the background scaling factor as: 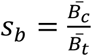, and the foreground scaling factor a 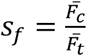.

### 2.7 Estimation of signal-alignment parameters *α* and *β*

Ryder estimates the signal-alignment parameters *α* and *β* using foreground signal values. Foreground regions are divided into a small number of bins, with 5 bins per region used by default for this fitting step. Values with zero signal in either sample are removed, and the remaining values are log_2_ transformed. The target foreground signal is first scaled by *s*_*f*_. Let *y*_*j*_ = log_2_(*s*_*f*_*t*_*j*_), *x*_*j*_ = log_2_(*c*_*j*_), where *j* indexes positive binned foreground signal values. In the default mode, Ryder aligns the target distribution to the reference distribution using a z-score-like distribution-matching transformation: 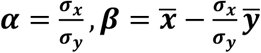, where ***σ*** indicates the standard deviation.

Ryder also provides an alternative linear fitting (lr) mode. In this mode, *α* and *β* are estimated by regressing the control/reference log_2_ signal *x*_*j*_ on the scaled target log2 signal *y*_*j*_, so that the transformed target value *αy*_*j*_ + *β* is aligned to *x*_*j*_. This provides an alternative when the default distribution-matching approximation is not appropriate.

### 2.8 Estimation of the foreground vs. background cutoff

Ryder classifies target-sample bigWig intervals as background or foreground signal using a cutoff *τ*. This cutoff is estimated from the target sample using the retained foreground reference regions and their matched background regions. Positive foreground and background signal values are log_2_ transformed, and the cutoff is defined as the midpoint between the mean foreground and background log_2_ signal levels: 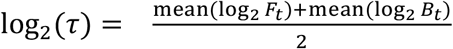, where *F*_*t*_ and *B*_*t*_ are positive target-sample signal values from foreground and background regions, respectively. Users may override the automatically estimated cutoff using the -noise option.

### 2.9 Detection of differential features with PATROL

After normalization, patrol.py can be used to quantify signal over user-defined genomic regions and identify differential features. PATROL merges the supplied regions into foreground intervals and generates background regions by shifting each foreground interval upstream and downstream by 10 region widths. These background regions are used to estimate a sample-specific noise level during signal quantification.

For each foreground region, PATROL computes the average bigWig signal in the control and treatment samples. If a region has zero average signal, the value is replaced by the estimated background noise level to avoid undefined log_2_ fold changes. PATROL then computes:

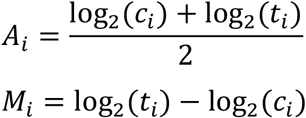

where *A*_*i*_is the average log_2_ signal and *M*_*i*_ is the log_2_ fold change for region *i*.

PATROL supports three differential-feature detection modes. In MD mode, it performs a two-pass Mahalanobis distance test on the (*M*_*i*_, *A*_*i*_) values. The first pass identifies outliers using all regions; the second pass recomputes the covariance matrix and center after excluding first-pass outliers, then recalculates distances and *p*-values for all regions. Regions with *p*-values below the user-defined cutoff and with average signal above the estimated noise threshold are reported as differential. In FC mode, PATROL combines a fold-change threshold with a Poisson test. The default fold-change threshold is | *M*_*i*_ |> 1, corresponding to a two-fold change, and the Poisson p-value cutoff is specified by -pcut. In the default FCn mode, PATROL uses the same fold-change threshold but does not apply the Poisson test. For all modes, regions with positive *M*_*i*_ are reported as treatment-enriched, whereas regions with negative *M*_*i*_ are reported as control-enriched. These modes are provided for different downstream use cases. The default FCn mode is intended for simple region-level screening and visualization using a fixed log2 fold-change threshold. The FC mode adds a Poisson-based filter when users want a count-like significance criterion in addition to fold change. The MD mode identifies regions with unusual M-A behavior relative to the overall region distribution. These options provide convenient exploratory feature detection from normalized signal tracks. PATROL outputs a statistics table, MA plots, BED files of differential regions, and aggregate signal plots for differential and total input regions.

PATROL is intended as a downstream feature-detection module for Ryder-normalized signal tracks. It provides region-level summaries using fold-change, Poisson-based, or Mahalanobis-distance-based criteria, but it does not explicitly model biological replicate variance. Ryder-normalized tracks can be used for visualization, aggregate-profile comparison, and exploratory region-level analysis. For formal differential analysis across biological replicates, Ryder can be applied to individual replicate bigWig tracks, and normalized signals or corresponding read counts can then be quantified per replicate and analyzed using established frameworks such as DESeq2 (Love, et al., 2014) or edgeR (Nikolayeva and Robinson, 2014). Thus, PATROL is not intended to replace replicate-aware differential peak-calling methods when biological replicate variation is the main inferential target.

### 2.10 Reference genome and sequencing data pre-processing

All analyses were performed using mouse (mm10) or human (hg38) reference genomes annotated by GENCODE (M21 and V30, respectively) (Frankish, et al., 2019). Reference genomes for Drosophila melanogaster (dm6) and Saccharomyces cerevisiae (sacCer3), used for spike-in normalization were obtained from the UCSC Genome Browser (Perez, et al., 2024). Published ChIP-seq and ATAC-seq, and DNase-seq datasets generated in this study were processed from raw reads into reads-per-million (RPM)-normalized bigWig tracks following established pipelines (Cao, et al., 2023; Cui, et al., 2023). Briefly, sequencing reads were mapped using Bowtie2 (v2.3.5) (Langmead and Salzberg, 2012), retaining only non-redundant reads with MAPQ ≥ 10 for downstream analyses. Peak calling was performed using the cLoops2 callPeaks module (Cao, et al., 2022) with pooled replicates and assay-specific parameters: DNase-seq (-eps 75,150 -minPts 10,20 -sen), ATAC-seq (-eps 75,150 -minPts 20,30 -sen), H3K4me3 ChIP-seq (-eps 150,300 -minPts 20,30 -sen), H3K27me3 ChIP-seq (-eps 300,500 -minPts 20,30 -sen), and H3K9ac ChIP-seq (-eps 75,150 -minPts 30,50 -sen, input as control). Genome-browser-style visualizations were generated using the cLoops2 plotting module (Cao, et al., 2022), and averaged tracks from biological replicates were produced using the bigwigCompare function from deepTools2 (Ramirez, et al., 2016).

### 2.11 Nucleosome center-weighted occupancy score

A revised nucleosome center-weighted occupancy score (Brogaard, et al., 2012) is computed for all nucleosome fragment reads (147 ± 15 *bp*) to quantify aggregated nucleosome density in MNase-seq data. Specifically, a normalized vector of Gaussian weights is generated to weight each nucleosome fragment based on its distance from the fragment center and on whether the fragment length is odd or even. The weight is defined 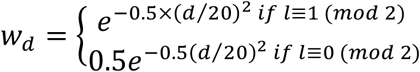, where *d* is the base distance from the fragment center and *l* is the fragment length. The resulting weight vector is normalized by dividing each element by the sum of all weights, ensuring that the total sums to 1. Finally, the genome-wide nucleosome center-weighted occupancy score is normalized by dividing by the total number of nucleosomes, allowing for comparisons between pseudo-bulk samples.

## 3 Results

### 3.1 Ryder normalizes epigenomic signal tracks using internal reference regions

We encountered the normalization problem that motivated Ryder development in a DNase-seq dataset comparing wild-type and *Gata3* knockout DN3 (CD4-CD8-CD25+) thymocytes. In this experiment, *Gata3*^fl/fl^-CreERT mice were treated with tamoxifen to induce *Gata3* deletion. *Gata3* knockout (KO) resulted in a markable shrunken thymus (**Figure S1A–C**), consistent with the essential role of GATA3 in thymocyte development that its deletion halted T cell development at subsequent stages (Pai, et al., 2003). We anticipated changes primarily in chromatin accessibility at GATA3-bound DNase hypersensitive sites (DSHs). However, after standard library-size normalization by reads per million (RPM), we observed a global increase in DNase-seq signal in the KO condition, including at the assumed invariant CTCF binding sites (**Figure 1A** and **B**). These sites were defined as DHSs that overlapped curated reference CTCF sites but did not overlap with GATA3 binding sites, which should not be changed by *Gata3* KO, thus could serve as the internal reference regions. Therefore, the global increase was likely attributable to technical variation and requiring proper normalization.

**Figure 1.**
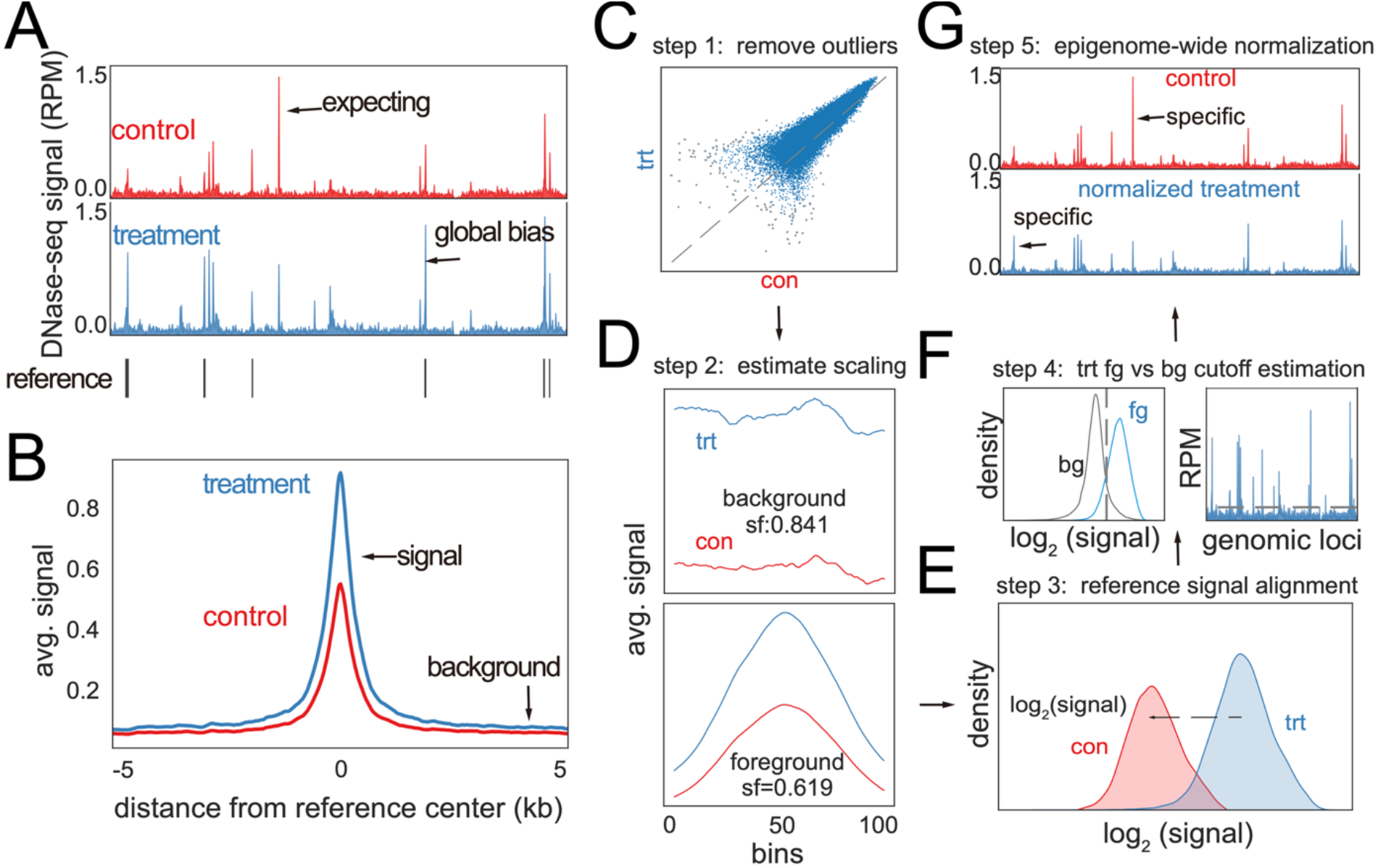
Ryder Normalization Workflow. (A) Genome browser view of DNase-seq signals in wild-type (control) versus GATA3 knockout (treatment) samples. The tracks represent the average signal from two biological replicates after normalization by library size. Reference peaks shown below the tracks are DNase hypersensitive sites (DHSs) from the control sample that overlap with invariant CTCF binding sites, defined using CTCF ChIP-seq peaks from Cistrome (L’Yi, et al., 2023) and data from more than 50 cell and tissue types sharing the CTCF sites (**Methods**). (B) Aggregate DNase-seq signals at reference regions (CTCF sites overlapping DNase-seq peaks) before normalization, illustrating the initial discrepancies in background and signal levels between WT and GATA3 KO samples. (C) Schematic of the Ryder normalization workflow step 1: identification and removal of outlier internal reference regions using Mahalanobis distance on M-A transformed log_2_ signals. (D) Schematic of the Ryder normalization workflow step 2: estimation of scaling factors for background and reference signal regions. (E) Schematic of the Ryder normalization workflow step 3: signal alignment via z-score transformation of the target sample to the control sample, or alternatively by fitting a linear model (log_2_ scale). (F) Schematic of the Ryder normalization workflow step 4: classification of genomic bins into background or signal based on a cutoff defined as the midpoint between the mean log_2_ foreground and background signal levels. (G) Schematic of the Ryder normalization workflow step 5: application of normalization across the epigenome using background- and signal-specific transformations.

The Ryder workflow is summarized in **Figure 1C-G** and described in detail in **Methods**. Briefly, candidate internal reference regions are first filtered to remove outliers, and the retained reference regions are used to define foreground regions and nearby background regions. Ryder estimates separate scaling factors for background and foreground regions, aligns the foreground signal distribution in log_2_ space, estimates a foreground/background cutoff, and applies the resulting two-tier transformation genome-wide (**Figure 1C–G**).

### 3.2 Ryder corrects technical signal inflation in *Gata3* knockout DNase-seq

Using invariant CTCF-associated DHSs as internal reference regions, Ryder aligned the reference-region signals between wild-type and KO samples (**Figure 2A and C**). After Ryder normalization, GATA3-bound DHSs showed significantly reduced accessibility in the knockout condition (**Figure 2B and D**), consistent with the expected role of GATA3 in maintaining accessibility at its bound regulatory elements. Thus, Ryder did not simply erase biological differences; instead, it corrected the reference-region bias and revealed the expected GATA3-dependent accessibility loss.

**Figure 2.**
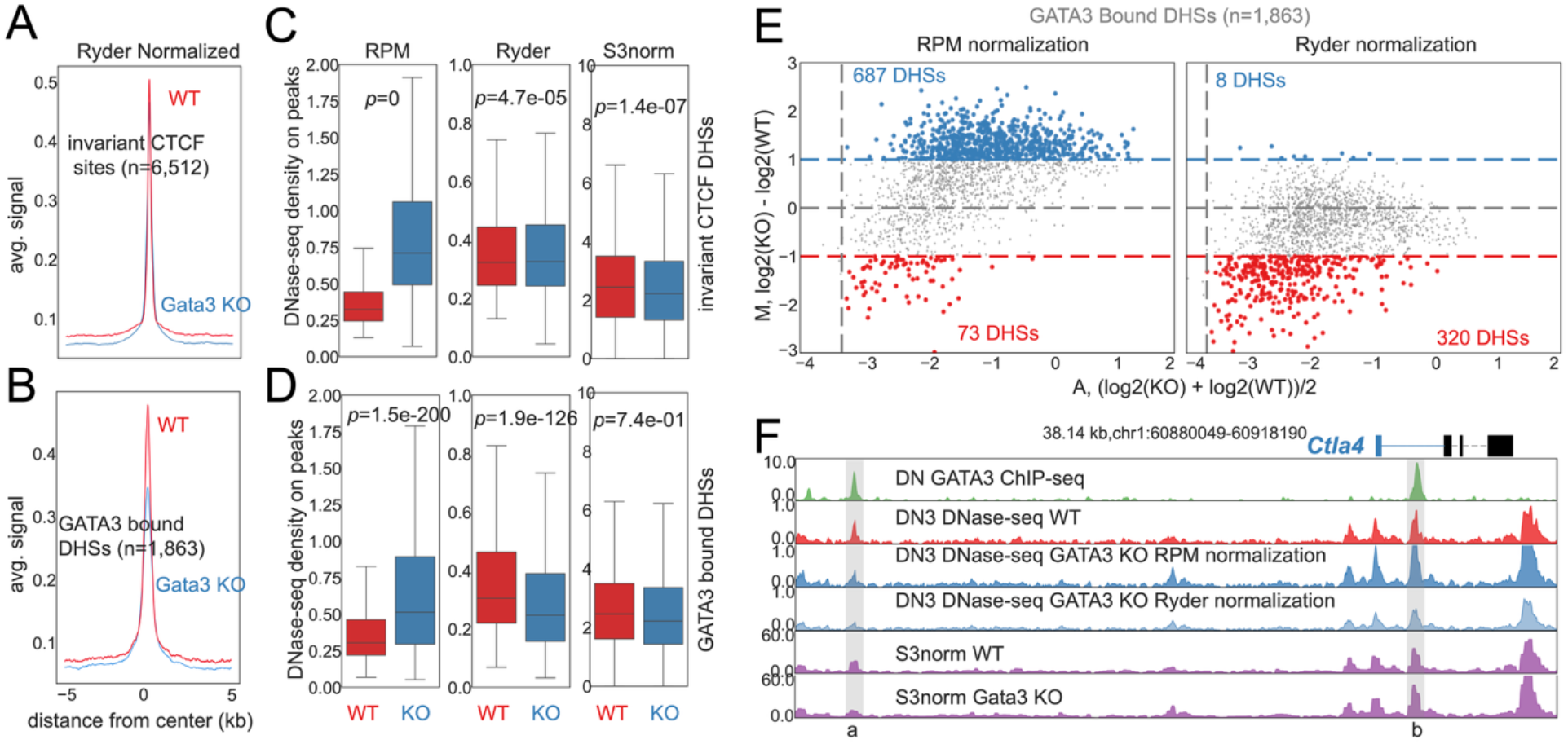
Ryder normalization reveals reduced chromatin accessibility at GATA3-bound DHSs after GATA3 knockout. (A) Aggregate Ryder-normalized DNase-seq signals centered on invariant CTCF-associated DNase hypersensitive sites (DHSs) in wild-type (WT) and Gata3 knockout (KO) DN3 thymocytes. These CTCF-associated DHSs were used as internal reference regions for Ryder normalization. (B) Aggregate Ryder-normalized DNase-seq signals centered on GATA3-bound DHSs, showing reduced chromatin accessibility in Gata3 KO cells compared with WT cells. (C) Boxplots comparing DNase-seq density at invariant CTCF-associated DHSs under RPM normalization, Ryder normalization and S3norm normalization. Invariant CTCF-associated DHSs were defined as DHSs that overlapped curated reference CTCF sites but did not do not overlap with GATA3 binding sites. Ryder substantially reduced the global signal difference between WT and Gata3 KO at the internal reference regions. P-values are indicated above each comparison and calculated across matched genomic regions using a two-sided paired Wilcoxon signed-rank test. (D) Boxplots comparing DNase-seq density at GATA3-bound DHSs under RPM normalization, Ryder normalization and S3norm normalization. After Ryder normalization, GATA3-bound DHSs retain significantly reduced accessibility in Gata3 KO cells, consistent with the expected role of GATA3 in maintaining accessibility at its bound regulatory elements. (E) MA-plots of significantly altered GATA3-bound DHSs (fold change ≥ 2) between mouse wild-type and GATA3 knockout double-negative (DN) thymocytes before (left panel) and after Ryder normalization (right panel). Normalization reveals additional differential DHSs that were not detected in the unnormalized data due to global differences. (F) Genome browser view of DNase-seq data from DN3 GATA3 KO cells before and after Ryder or S3norm normalization. Two highlighted GATA3-bound DHSs, undetected prior to normalization, become apparent after normalization, demonstrating the method’s ability to uncover biologically relevant changes.

We further compared RPM and Ryder normalization using MA plots of GATA3-bound DHSs. Under RPM normalization, many GATA3-bound DHSs appeared increased in the knockout sample, consistent with the global signal inflation. After Ryder normalization, the number of KO-increased GATA3-bound DHSs was reduced from 687 to 8, whereas the number of KO-decreased GATA3-bound DHSs increased from 73 to 320 (**Figure 2E**). This effect is exemplified by two enhancers bound by GATA3 in the *Ctla4* gene (**Figure 2F**, marked as a & b). *Ctla4* encodes an important immune checkpoint protein (Buchbinder and Desai, 2016), and its expression can be decreased by GATA3 KO, further affecting the downstream T cell development.

By comparison, S3norm (Xiang, et al., 2020), a similar signal-track normalization method, did not address the normalization bias in this dataset. After S3norm normalization, invariant CTCF-associated DHSs have significantly lower signals in KO samples (**Figure 2C** and **Figure S1D**). Moreover, S3norm produced problematic signal patterns in the GATA3-bound DHSs surrounding background regions (**Figure S1E**) and failed to detect differences in signal density at GATA3-bound DHSs (**Figure 2D & F**). Additionally, Ryder was computationally more efficient than S3norm and completed normalization in less time and consumed less memory (**Figure S1F**). Together, these results show that Ryder provides a biologically interpretable correction for the DNase-seq dataset while requiring less runtime and memory than S3norm in this benchmark.

### 3.3 Stable TSSs can serve as alternative internal reference regions

Although invariant CTCF sites were used as the default reference set in most analyses, Ryder is not restricted to CTCF-based references. To test an alternative reference strategy, we analyzed a public CD8+ T cell ATAC-seq dataset comparing resting and stimulated cells, for which matched RNA-seq data allowed us to identify genes with stable expression. This dataset was previously used in IGN (Hu, et al., 2025) to demonstrate the algorithm. Based on RNA-seq, we selected 500 conditionally stable genes with the smallest fold changes and an expression level greater than 1 RPKM in both conditions. ATAC-seq peaks overlapping TSSs of these genes were then used as an alternative internal reference set (**Figure S2A**). For comparison, we also defined a stable CTCF reference set by selecting ATAC-seq peaks shared between the two conditions that further overlapped curated invariant CTCF sites.

After Ryder normalization, both the stable TSS and CTCF reference sets showed well-aligned reference signals while preserving the global activation-associated increase in chromatin accessibility (**Figure S2B**). In particular, ATAC-seq signal significantly increased at TSSs of upregulated genes after stimulation with both reference sets (**Figure S2C**), whereas TSSs of downregulated genes did not show an inappropriate global increase after Ryder normalization with CTCF-based reference (**Figure S2C**). IFN-γ is a critical effector cytokine and a well-known functional marker of CD8+ T cell activation (Ghanekar, et al., 2001), and its expression is tightly controlled by *cis*-regulatory elements (Cui, et al., 2023). At the *Ifng* locus, the original RPM-normalized ATAC-seq tracks showed an apparent decrease in promoter and nearby enhancers accessibility after stimulation (region b in **Figure S2D**), inconsistent with the highly upregulated gene expression. After Ryder normalization using either CTCF- or stable TSS-based reference sets, stimulation-associated accessibility was preserved at the *Ifng* promoter and nearby regulatory regions. Among these, region a showed a prominent de novo gain of accessibility after stimulation and represents a putative enhancer candidate that may contribute to activation-induced *Ifng* expression (**Figure S2D**). Thus, Ryder normalization makes the ATAC-seq profile at the *Ifng* locus more consistent with the strong RNA-seq induction observed after CD8 T cell stimulation.

These results demonstrate that Ryder can use biologically justified alternative reference sets, such as stable TSSs, when CTCF is not appropriate for a given perturbation.

### 3.4 DNase-seq analysis of BRG1-AID cells with spike-in controls reveals limitations of single-factor normalization

BRG1 is a core chromatin remodeler required for maintaining enhancer accessibility (Hagihara, et al., 2025; King and Klose, 2017; Ren, et al., 2024). Therefore, we expected that BRG1 depletion reduces chromatin accessibility preferentially at BRG1-bound enhancers, with the promoter accessibility less affected.

We further evaluated Ryder’s performance by applying it to DNase-seq experiments comparing wild-type cells to those with acute BRG1 depletion using the auxin-inducible degron (AID) system in mouse fibroblast cells, incorporating human 293T cells as spike-in controls (**Figure 3A**). This experimental design mirrors that of our previous study (Ren, et al., 2024), which showed that BRG1 plays a critical role in maintaining the accessibility of chromatin at enhancers. Clear DHSs were observed around the *GAPDH* locus in both human spike-in cells and mouse samples (**Figure S3A**), along with enriched signal profiles (**Figure S3B**).

**Figure 3.**
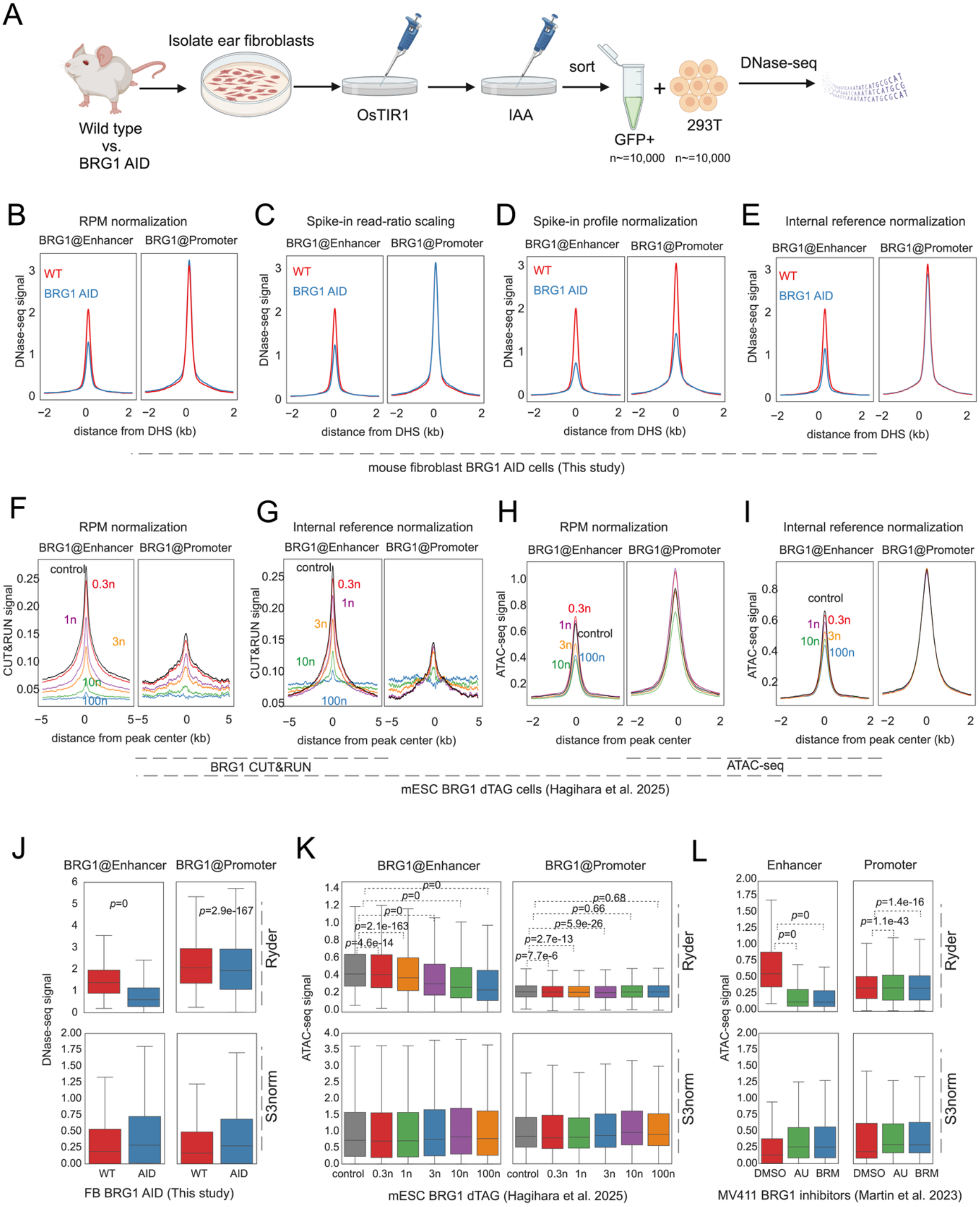
Comparison of spike-in, S3norm, and internal-reference normalization in BRG1 perturbation datasets. (A) Schematic of the DNase-seq spike-in experiment in mouse fibroblasts. Ear fibroblasts were isolated from wild-type and BRG1-AID mice, transfected with OsTIR1, treated with IAA to induce BRG1 degradation, and sorted for GFP-positive cells. Approximately equal numbers of mouse fibroblasts and human 293T spike-in cells were mixed before DNase-seq library preparation. (B) Aggregate DNase-seq signals at DHSs in mouse fibroblasts, comparing wild-type and BRG1-AID samples. DHSs were categorized into BRG1-bound enhancers and promoters (within ±2 kb of TSS). Signals were normalized using reads per million (RPM). BRG1 binding sites were identified from ChIC-seq data generated in a previous study (Ren, et al., 2024) and overlapped with DHSs called from wild-type DNase-seq data generated in this study. (C) Same as (B), but signals further scaled by the ratio of spike-in read counts between samples (Orlando, et al., 2014). (D) Same as (B), with additional normalization using spike-in signal profiles: human 293T DHSs normalized to derive parameters applied to mouse data. (E) Same as (B), with further normalization using internal reference regions—mouse DHSs overlapping invariant CTCF sites and not overlapping BRG1 peaks. (F) Aggregate BRG1 CUT&RUN profiles in mouse embryonic stem cells (mESCs) following a titration of the dTAG13 degrader, which induces varying levels of BRG1 protein degradation. Signals were RPM-normalized and plotted separately for BRG1-bound enhancers and promoters (as defined in panel B), based on overlap with ATAC-seq data from the same study. Data were obtained from (Hagihara, et al., 2025), see details in Supplemental Information. (G) Same as (F), but with CUT&RUN signals normalized using Ryder’s internal reference method. The reference set consists of invariant CTCF sites that do not overlap BRG1-bound peaks. (H) Same as (F), but showing the corresponding ATAC-seq profiles from the same dTAG13 degrader titration, illustrating the impact of BRG1 depletion on chromatin accessibility. (I) Same as (H), but with ATAC-seq signals normalized using Ryder’s internal reference method. (J) Boxplots comparing Ryder-normalized and S3norm-normalized DNase-seq signals at BRG1-bound enhancers and promoters in wild-type and BRG1-AID mouse fibroblasts. The upper panels show Ryder-normalized signals, and the lower panels show S3norm-normalized signals. (K) Boxplots comparing Ryder-normalized and S3norm-normalized ATAC-seq signals at BRG1-bound enhancers and promoters across the mESC BRG1 dTAG13 titration series. The upper panels show Ryder-normalized signals, and the lower panels show S3norm-normalized signals. Dashed brackets indicate pairwise comparisons with the control condition. (L) Boxplots comparing Ryder-normalized and S3norm-normalized ATAC-seq signals at enhancers and promoters in MV411 cells treated with DMSO and BRG1 inhibitor AU15330 or BRM014. The upper panels show Ryder-normalized signals, and the lower panels show S3norm-normalized signals. AU, AU15330; BRM, BRM014. For boxplots in (J–L), boxes indicate the interquartile range, center lines indicate medians, and whiskers indicate the distribution range excluding outliers. P-values shown above comparisons were calculated across matched genomic regions using a two-sided paired Wilcoxon signed-rank test.

To assess the effects of BRG1 depletion, we compared DNase-seq signals at BRG1-bound enhancers and promoters using normalization by reads per million (RPM) first (**Figure 3B**), then incorporating the ratio of spike-in read counts (Orlando, et al., 2014) for additional correction (**Figure 3C**). Ryder further modeled background and signal-specific scale factors (*s*_*b*_,*s*_*f*_) and signal alignment parameters (***α, β***), using either all spike-in DHSs (**Figure 3D**) or DHSs overlapping invariant CTCF binding sites as internal reference (**Figure 3E**). All methods consistently revealed reduced accessibility at BRG1-bound enhancers after BRG1 depletion (**Figure S3C&D**), but internal reference-based normalization identified more significantly decreased sites at both enhancers and promoters than RPM or spike-in ratio normalizations (**Figure S3D**). Additionally, in the absence of spike-in controls, S3norm failed to recover the expected enhancer-preferential reduction in accessibility after BRG1 depletion and instead produced globally inflated signal distributions (**Figure 3J**).

These results show that spike-in controls are useful but not automatically definitive: different spike-in-based strategies can produce different corrections, whereas Ryder provides an internal reference framework whose stability assumption can be evaluated directly in the endogenous genome.

### 3.5 Ryder recovers BRG1-dependent enhancer changes across CUT&RUN and ATAC-seq perturbation datasets

To test whether Ryder generalizes beyond our generated DNase-seq data, we reanalyzed a recent dataset that used the dTAG system for progressive degradation of BRG1, followed by CUT&RUN and ATAC-seq (Hagihara, et al., 2025).

Although simple RPM normalization of the CUT&RUN data suggests a dose-dependent decrease in BRG1 binding, this interpretation is confounded by a concurrent artifactual shift in the background signal (**Figure 3F**). In contrast, Ryder normalization corrects for this background noise and confirms a clear, dose-dependent loss of BRG1 binding specifically at enhancers, with minimal changes at promoters except with extensive degradation of BRG1 (**Figure 3G**). Strikingly, Ryder also uncovers the functional consequence of this binding loss: a corresponding, dose-dependent decrease in chromatin accessibility at the enhancer regions (**Figure 3I**). This subtle but significant trend, which was not apparent with simpler methods (**Figure 3H**), is consistent with our BRG1-AID results and reinforces BRG1’s direct role in maintaining enhancer accessibility. By comparison, S3norm did not resolve a comparable enhancer-specific dose-response pattern (**Figure 3K**). Together, these analyses show that by leveraging an internal reference, Ryder provides the required sensitivity to detect these quantitative biological trends, meanwhile highlighting the risks of relying on spike-in controls that may not be suitable for every assay.

Based on the expectation that BRG1 primarily affects enhancer accessibility, we further evaluated Ryder on a published independent ATAC-seq dataset from human MV411 cells treated with BRG1 inhibitors and with *Drosophila* S2 cells as spike-in controls (Martin, et al., 2023). Clear ATAC-seq peaks were observed around the *GAPDH* locus in MV411 cells but not spike-in S2 cells whereas the displayed *Drosophila* spike-in locus showed a distinct and less comparable local signal pattern (**Figure S3E**). Nevertheless, aggregate profiles confirmed that S2 spike-in cells had enriched ATAC-seq signal at their own peak regions (**Figure S3F**). These observations suggest that, although the spike-in data contained usable accessibility signal, the spike-in and target-cell chromatin profiles were not necessarily equivalent. As expected, Ryder’s internal reference normalization robustly confirmed a significant reduction in enhancer accessibility, while promoter accessibility remained largely unchanged (**Figure S3G&K**). Critically, this stands in contrast to standard RPM normalization, which produced a misleading, artifactual increase in promoter accessibility (**Figure S3H**). By avoiding this common artifact, Ryder provides a more accurate representation of the underlying biology, highlighting the importance of a robust normalization strategy. While spike-in-based methods are available, they can also introduce unwanted bias if not carefully implemented, as shown by the variable results from ratio-based versus profile-based approaches (**Figure S3I&J**). Quantitative analysis further confirmed that Ryder detected decreased enhancer accessibility after AU15330 and BRM014 treatment, whereas promoter accessibility was much less affected (**Figure 3L**). By comparison, S3norm produced weaker and less biologically interpretable enhancer–promoter distinctions in this dataset (**Figure 3L**).

Together, these analyses show that Ryder can recover BRG1-dependent enhancer accessibility changes across DNase-seq, CUT&RUN, and ATAC-seq datasets, while avoiding artifacts associated with single-factor or poorly calibrated spike-in normalization.

### 3.6 Ryder preserves expected global changes in MNase-seq and ChIP-seq

Additionally, we demonstrate that Ryder can normalize nucleosome data by applying a background scaling factor *s*_*b*_, estimated from MNase-seq signals at invariant CTCF sites (**Figure S4A**). Using this normalization approach, we found that inhibition of BRG1 using a chemical inhibitor, BRM014, resulted in increased nucleosome occupancy at enhancer centers and decreased occupancy in flanking regions, consistent with our previous finding that BRG1 acts to maintain a nucleosome-free region at enhancers (Hu, et al., 2011; Ren, et al., 2024). At promoters, the density of the +1 nucleosome increased by the inhibitor treatment, whereas other nucleosomes near the TSS showed subtle changes (**Figure 4A**). Because the nucleosome-to-background signal difference is modest, normalization using only the background scaling factor effectively removes global background variations, yielding results consistent with chromatin accessibility patterns observed by DNase-seq (**Figure 3E**) and ATAC-seq (**Figure 3I** and **Figure S3K**).

**Figure 4.**
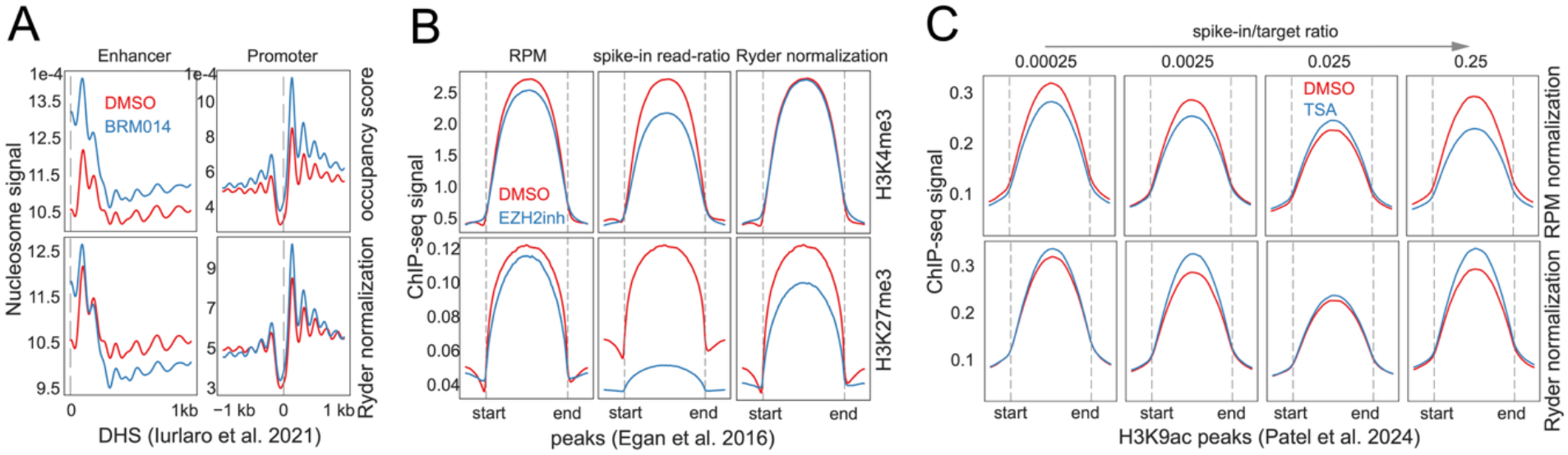
Ryder normalization preserves expected nucleosome and histone-modification changes in MNase-seq and ChIP-seq datasets. (A) Aggregate MNase-seq nucleosome occupancy profiles in mESCs treated with DMSO or BRM and BRG1 inhibitor BRM014, separated into enhancers and promoters. Upper panel: nucleosome occupancy score (**Methods**). Lower panel: signals further normalized using internal reference regions—DHSs overlapping invariant CTCF sites without BRG1 binding. MNase-seq data were reprocessed from (Iurlaro, et al., 2021), see details in Supplemental Information. (B) Aggregate ChIP-seq signals in human PC9 cells for H3K4me3 (top) and H3K27me3 (bottom), treated with DMSO or EZH2 inhibitor (GSK126). Columns show normalization by: (1) RPM; (2) spike-in read count ratios; (3) internal reference peaks overlapping invariant CTCF sites. Data were obtained from (Egan, et al., 2016), see details in Supplemental Information. (C) Aggregated H3K9ac ChIP-seq signals in HeLa-S3 cells treated with DMSO or Trichostatin A (TSA), which is HDACs inhibitor, with variable yeast spike-in ratios. Upper panel: RPM-normalized signals. Lower panel: normalized using peaks overlapping invariant human CTCF sites. Data were obtained from (Patel, et al., 2024), see details in Supplemental Information.

Finally, we validated our internal reference normalization method using ChIP-seq datasets in which global changes in chromatin modifications are well-established (Egan, et al., 2016; Patel, et al., 2024). These scenarios are critical tests for any normalization strategy. First, cells treated with an inhibitor of EZH2, the catalytic subunit of the PRC2 H3K27 methyltransferase, are expected to exhibit a global decrease in H3K27me3 while maintaining stable H3K4me3 levels; Ryder accurately reflected this outcome (**Figure 4B**). Our method correctly quantified the reduction in H3K27me3 without introducing the common artifact of an apparent increase in the stable H3K4me3 mark, a problem often seen with RPM normalizations (**Figure 4B & Figure S4B**). Second, Ryder confirmed the anticipated global increase in histone H3K9ac levels in cells treated with HDAC inhibitors (**Figure 4C**). The success in both of these distinct and challenging scenarios demonstrates Ryder’s utility for accurately analyzing ChIP-seq data, particularly when global epigenetic shifts are expected.

## 4 Conclusion and Discussion

Accurate normalization is critical for interpreting epigenomic data, yet many existing methods are often limited by untestable assumptions or inflexibility in the face of global chromatin changes. Here, we present Ryder, a Python tool that overcomes these challenges by using a flexible, two-tier normalization strategy anchored by stable internal reference regions. This approach separately corrects for background noise and signal intensity to achieve consistent signal-to-noise ratios across samples, providing a more robust correction than single-factor methods. Ryder also empowers users to select a single-factor normalization, derived from either signal or background, offering a simpler alternative for specific use cases.

By leveraging stable internal reference regions, such as invariant CTCF sites, Ryder aims to effectively distinguish genuine biological signals from technical artifacts. It supports a variety of epigenomic assays, including DNase-seq, CUT&RUN, ATAC-seq, MNase-seq, and ChIP-seq, with or without spike-in controls. Our validation studies demonstrate Ryder’s ability to accurately detect diverse biological changes, such as decreased accessibility at critical GATA3-bound enhancers following *Gata3* deletion, reduced enhancer accessibility following BRG1 degradation or inhibition, and expected chromatin modification changes following EZH2 and HDAC inhibitor treatments. We believe Ryder’s flexibility and accuracy can significantly enhance the reliability and interpretability of epigenomic analyses.

While spike-in controls can be useful for normalization, their implementation poses significant challenges regarding selection, titration, and the assumption that they faithfully mimic endogenous chromatin dynamics, which often remains unverified. Our re-analysis of the BRG1-dTAG dataset (Hagihara, et al., 2025) serves as a critical case study. The authors themselves concluded that their *Drosophila* spike-in was unusable due to inconsistent application. This highlights a fundamental advantage of internal references: they are inherently subjected to the exact same experimental conditions as the regions of interest. Furthermore, normalization strategies using spike-ins also vary, from applying a single global scaling factor to aligning specific spike-in peaks. As our analysis of the BRG1-degradation DNase-seq and BRG1-inhibition ATAC-seq data show, spike-in normalization can produce variable and potentially biased results depending on the strategy chosen. In contrast, an internal reference approach provides a more consistent and arguably safer baseline because its core assumption—the stability of specific genomic sites—is transparent and can be readily tested within the dataset itself. Ryder offers what we hope is a practical alternative by utilizing internal reference regions with readily testable assumptions, while still retaining the flexibility to integrate spike-ins when their use is demonstrably beneficial.

Ryder is not designed for direct normalization of individual sparse single-cell epigenomic profiles. It can be applied to pseudo-bulk single-cell epigenomic data after cells are aggregated by cell type, cluster, or treatment condition. This is useful when single-cell datasets are collected across multiple batches and then clustered into biologically meaningful groups, or when control and treatment pseudo-bulk profiles differ in signal-to-noise ratio. In this setting, aggregated fragments can be converted into RPM-normalized bigWig tracks and normalized by Ryder using stable reference regions with sufficient coverage. Because low coverage can make foreground and background parameter estimates unstable, users should apply Ryder only to pseudo-bulk groups with adequate signal and inspect Ryder diagnostic plots before interpretation.

Ryder addresses a different problem from count-matrix normalization methods such as DESeq2-style median-ratio normalization. Median-ratio approaches are valuable for replicate-aware differential testing after reads have been counted over genes, peaks, or other predefined regions, but they estimate a single sample-level scaling factor and do not generate genome-wide normalized signal tracks. In contrast, Ryder is designed for signal-track normalization when background noise, stable reference-region signal, and foreground signal intensity may shift differently between samples. Thus, Ryder is complementary to count-based statistical frameworks: Ryder can generate normalized bigWig tracks for visualization and signal-track comparison, and replicate-level Ryder-normalized signals or counts can subsequently be analyzed with replicate-aware methods when formal statistical inference is required.

In conclusion, Ryder offers a versatile and reliable solution for the normalization of diverse epigenomic data. By effectively distinguishing biological signal from technical noise, it significantly enhances the accuracy and interpretability of epigenomic analyses.

## Data and code availability

The DNase-seq datasets generated in this study, including wild-type versus Gata3 knockout DN3 thymocytes and control versus BRG1-AID fibroblasts with human 293T spike-in cells, have been deposited in the Gene Expression Omnibus database with accession number: **GSE300647**. Public datasets reanalyzed in this study are listed in Supplementary Table S6 with accession numbers. Ryder source code, documentation, test data and preprocessed reference-region files are available at https://github.com/YaqiangCao/ryder. Essential datasets and code to reproduce the results presented in this manuscript are available at: https://github.com/YaqiangCao/ryder/tree/main/demo. The exact software version used for the analyses in this manuscript has been archived on Zenodo under DOI: 10.5281/zenodo.21267457.

## Authors Contributions

Y.C. conceived the idea, designed the software package, and performed data analysis.

G.G. conducted the experiments. Y.C., G.G., and K.Z. co-wrote the manuscript. K.Z. supervised the study.

The authors declare no competing financial interests.

## Acknowledgements

This research was supported by the Intramural Research Program of the National Institutes of Health (NIH). The contributions of the NIH author(s) were made as part of their official duties as NIH federal employees, are in compliance with agency policy requirements, and are considered Works of the United States Government. However, the findings and conclusions presented in this paper are those of the author(s) and do not necessarily reflect the views of the NIH or the U.S. Department of Health and Human Services.

We thank the NHLBI DNA Sequencing Core Facility for sequencing the libraries and the NHLBI Flow Cytometry Core facility for sorting cells. This work utilized the computational resources of the NIH HPC Biowulf cluster (http://hpc.nih.gov). This manuscript benefits from the NHLBI Chat (https://ai.nhlbi.nih.gov/chat), Enterprise Google Gemini, and Enterprise ChatGPT for writing polishing.

