## Supplemental Information for "Ryder: Epigenome normalization using a two-tier model and internal reference regions"

#### Materials and Methods

##### Mice

The BRG1-AID mice were generated and maintained in our lab as described previously (Ren, et al., 2024). The *Gata3*$^{fl/fl}$-CreERT mice were kindly provided by Jinfang Zhu in NIAID/NIH. Mice were bred and maintained in the NHLBI animal facilities. All the animal experiments were performed under a protocol approved by the NHLBI Animal Care and Use Committee.

##### The GATA3 knockout with tamoxifen treatment in DN3 T cells

Tamoxifen was administered to *Gata3*$^{fl/fl}$-CreERT mice by intraperitoneal injection three times (once per day, 2 mg tamoxifen in 150μl corn oil per injection). DN3 T cells were isolated from the thymus 3.5 days later after first tamoxifen injection. To sort DN3 T cells, CD4/CD8 T cells were labeled with anti-CD4/CD8 antibodies and removed using BioMag Goat anti-rat IgG beads from QIAGEN. The remaining cells were stained with linage markers (anti-CD4, CD8a, TCR gamma/delta, CD19, B220, Gr-1, CD11b, CD11c, Nk1.1 and Ter119) and anti-CD3, CD44, CD25. DN3 T cells were sorted as linage$^-$CD3$^-$CD44$^-$CD25$^+$.

##### The auxin induced BRG1 deletion in mice primary fibroblasts

Primary fibroblasts were isolated and cultured as described previously. In brief, the fibroblasts were isolated from ear in wild type or BRG1-AID mice and then maintained in DMEM medium (high glucose) containing 10% fetal bovine serum. The osTIR1-GFP plasmids were transfected into fibroblasts by Lipofectamine LTX (Cat. No. 15338100) and FuGENE HD (Cat. No. E2311) with the ratio of 1:1. The auxin was added to cells 24 h later after transfection and incubated for 8 ~ 10 h. GFP+ cells were sorted by flow cytometry for further experiments.

##### DNase-seq

DNase-seq were performed as previously described with minor modifications. Briefly, 10K live cells (mice fibroblasts) were sorted and fixed in 1% formaldehyde. The equal amount of fixed HEK293T cells were added as spike in control. The fixed cells were permeabilized and the DNase1 (home-made) were added to digest cells at 37 °C for 1 min. Reactions were stopped by adding EDTA, DTT and DNase1 were heat denatured at 55 °C for 10min. After reverse crosslinking, the DNA was purified and DNase-seq libraries were prepared using NEBNext® Ultra™ II DNA Library Prep Kit for Illumina.

Table S1. Mapping Metrics for DNase-seq Samples Generated in This Study

| map to mouse genome | TotalReads | MappingRatio(%) | totalMappedPETs (MAPQ>=10) | uniquePETs | redundancy | tssEnrichmentScore |
|---|---|---|---|---|---|---|
| DNase-seq_DN3_GATA3_WT_rep1 | 21367057 | 82.51 | 15477761 | 13656214 | 0.12 | 3.10 |
| DNase-seq_DN3_GATA3_WT_rep2 | 25907927 | 79.1 | 17998897 | 15636321 | 0.13 | 3.05 |
| DNase-seq_DN3_GATA3_KO_rep1 | 19260022 | 82.52 | 14340692 | 12294587 | 0.14 | 6.32 |
| DNase-seq_DN3_GATA3_KO_rep2 | 16714976 | 81.29 | 12246634 | 10651694 | 0.13 | 6.16 |
| DNase-seq_FB_BRG1_WT_rep1 | 59804001 | 17.74 | 9793516 | 4065237 | 0.58 | 15.39 |
| DNase-seq_FB_BRG1_WT_rep2 | 80586744 | 16.36 | 12183302 | 3756305 | 0.69 | 13.46 |
| DNase-seq_FB_BRG1_AID_rep1 | 40557941 | 21.79 | 8007511 | 6288068 | 0.21 | 15.36 |
| DNase-seq_FB_BRG1_AID_rep2 | 65310477 | 20.24 | 12183822 | 8743209 | 0.28 | 18.24 |
| map to human (spike-in) | | | | | | |
| DNase-seq_FB_BRG1_WT_rep1 | 59804001 | 28.22 | 15265034 | 6741276 | 0.56 | 6.51 |
| DNase-seq_FB_BRG1_WT_rep2 | 80586744 | 34.25 | 24937056 | 8151174 | 0.67 | 5.44 |
| DNase-seq_FB_BRG1_AID_rep1 | 40557941 | 17.5 | 6332945 | 4844088 | 0.24 | 9.13 |
| DNase-seq_FB_BRG1_AID_rep2 | 65310477 | 16.74 | 9774468 | 6826005 | 0.30 | 11.86 |

Table S2. Mapping Metrics for mESC BRG1 dTAG CUT&RUNG and ATAC-seq Samples from (Hagihara, et al., 2025)

| CUT&RUNG | | | | | | | |
|---|---|---|---|---|---|---|---|
| | SampleInformation | TotalReads | Mapping Ratio(%) | totalMappedReads | uniqueReads | redundancy | tssEnrichmentScore |
| GSM8447862 | Control CUT&RUNG rep1 | 40080542 | 71.82 | 47939904 | 33552797 | 0.30 | 2.35 |
| GSM8447863 | Control CUT&RUNG rep2 | 35173549 | 81.19 | 49665062 | 36344419 | 0.27 | 1.96 |
| GSM8447864 | 0.3n dTag CUT&RUNG rep1 | 34256209 | 74.9 | 43323418 | 30952016 | 0.29 | 1.97 |
| GSM8447865 | 0.3n dTag CUT&RUNG rep2 | 33071167 | 75.04 | 42881668 | 31076579 | 0.28 | 1.84 |
| GSM8447866 | 1n dTag CUT&RUNG rep1 | 36274468 | 78.69 | 47671180 | 34005191 | 0.29 | 1.80 |
| GSM8447867 | 1n dTag CUT&RUNG rep2 | 37401223 | 75.5 | 47772276 | 33483606 | 0.30 | 1.74 |
| GSM8447868 | 3n dTag CUT&RUNG rep1 | 35439068 | 82.74 | 48575338 | 35402699 | 0.27 | 1.38 |
| GSM8447869 | 3n dTag CUT&RUNG rep2 | 31777496 | 76.42 | 40941186 | 28812074 | 0.30 | 1.40 |

| | | | | | | | |
|---|---|---|---|---|---|---|---|
| GSM8447870 | 10n dTag CUT&RUNG rep1 | 37021905 | 81.41 | 49933356 | 35397011 | 0.29 | 1.13 |
| GSM8447871 | 10n dTag CUT&RUNG rep2 | 34743819 | 79.91 | 46275358 | 31385959 | 0.32 | 1.04 |
| GSM8447872 | 100n dTag CUT&RUNG rep1 | 35586795 | 88.04 | 52079120 | 39105705 | 0.25 | 1.03 |
| GSM8447873 | 100n dTag CUT&RUNG rep2 | 41295578 | 79.04 | 53828950 | 36112012 | 0.33 | 1.03 |
| ATAC-seq | | | | | | | |
| GSM8447848 | WT ATAC rep1 | 47596196 | 28.65 | 24264288 | 14586184 | 0.40 | 8.46 |
| GSM8447849 | WT ATAC rep2 | 32541134 | 26.11 | 15059296 | 9775762 | 0.35 | 8.45 |
| GSM8447850 | Control ATAC rep1 | 104414637 | 29.87 | 55392112 | 26424529 | 0.52 | 6.74 |
| GSM8447851 | Control ATAC rep2 | 38847627 | 24.05 | 16524674 | 10317716 | 0.38 | 8.61 |
| GSM8447852 | 0.3n dTag ATAC rep1 | 82653938 | 23.51 | 34664040 | 17327490 | 0.50 | 8.74 |
| GSM8447853 | 0.3n dTag ATAC rep2 | 85332832 | 26.99 | 41107056 | 20430218 | 0.50 | 8.31 |
| GSM8447854 | 1n dTag ATAC rep1 | 110718100 | 24.31 | 47791140 | 21409516 | 0.55 | 8.00 |
| GSM8447855 | 1n dTag ATAC rep2 | 89200833 | 21.58 | 33823302 | 15398627 | 0.54 | 10.06 |
| GSM8447856 | 3n dTag ATAC rep1 | 111904590 | 25.8 | 51308270 | 23258592 | 0.55 | 7.60 |
| GSM8447857 | 3n dTag ATAC rep2 | 86297162 | 24.94 | 38026776 | 18324483 | 0.52 | 8.12 |
| GSM8447858 | 10n dTag ATAC rep1 | 104749972 | 30.85 | 57610440 | 28697349 | 0.50 | 5.85 |
| GSM8447859 | 10n dTag ATAC rep2 | 92679211 | 25.55 | 41835884 | 20177157 | 0.52 | 7.94 |
| GSM8447860 | 100n dTag ATAC rep1 | 105471551 | 29.19 | 54765438 | 26688772 | 0.51 | 6.24 |
| GSM8447861 | 100n dTag ATAC rep2 | 89052066 | 21.34 | 33440832 | 15154868 | 0.55 | 10.00 |

#### Table S3. Mapping Metrics for MV411 BRG1 Inhibitors Treated ATAC-seq Samples from (Martin, et al., 2023)

| map to fly (spike-in) | SampleInformation | TotalReads | MappingRatio(%) | totalMappedReads | uniqueReads | redundancy | tssEnrichmentScore |
|---|---|---|---|---|---|---|---|
| GSM7695921 | MV411_ATACseq_1h_AU15330_rep1 | 17422822 | 2.91 | 798838 | 579152 | 0.28 | 2.05 |
| GSM7695922 | MV411_ATACseq_1h_AU15330_rep2 | 17463612 | 2.85 | 781684 | 562673 | 0.28 | 1.92 |
| GSM7695923 | MV411_ATACseq_1h_AU15330_rep3 | 17030548 | 3.22 | 860174 | 583944 | 0.32 | 1.95 |
| GSM7695924 | MV411_ATACseq_1h_BRM014_rep1 | 17983974 | 2.86 | 812070 | 584247 | 0.28 | 2.07 |
| GSM7695925 | MV411_ATACseq_1h_BRM014_rep2 | 17557774 | 3.32 | 918844 | 610007 | 0.34 | 1.93 |
| GSM7695926 | MV411_ATACseq_1h_BRM014_rep3 | 19013075 | 2.73 | 813036 | 580961 | 0.29 | 1.99 |
| GSM7695927 | MV411_ATACseq_1h_DMSO_rep1 | 11644740 | 1.88 | 351320 | 243360 | 0.31 | 1.96 |
| GSM7695928 | MV411_ATACseq_1h_DMSO_rep2 | 15019014 | 1.86 | 438878 | 325268 | 0.26 | 1.93 |
| GSM7695929 | MV411_ATACseq_1h_DMSO_rep3 | 15889335 | 1.68 | 416380 | 310511 | 0.25 | 1.87 |
| map to human | | | | | | | |
| GSM7695921 | MV411_ATACseq_1h_AU15330_rep1 | 17422822 | 90.38 | 27910554 | 18754660 | 0.33 | 14.51 |
| GSM7695922 | MV411_ATACseq_1h_AU15330_rep2 | 17463612 | 90.75 | 28238916 | 19480882 | 0.31 | 14.34 |
| GSM7695923 | MV411_ATACseq_1h_AU15330_rep3 | 17030548 | 91.99 | 27851284 | 18909219 | 0.32 | 15.41 |
| GSM7695924 | MV411_ATACseq_1h_BRM014_rep1 | 17983974 | 90.23 | 28708372 | 19500522 | 0.32 | 13.68 |
| GSM7695925 | MV411_ATACseq_1h_BRM014_rep2 | 17557774 | 91.68 | 28502348 | 19443913 | 0.32 | 14.60 |
| GSM7695926 | MV411_ATACseq_1h_BRM014_rep3 | 19013075 | 90.48 | 30731340 | 22023101 | 0.28 | 12.76 |
| GSM7695927 | MV411_ATACseq_1h_DMSO_rep1 | 11644740 | 94.22 | 19879902 | 14074214 | 0.29 | 14.43 |
| GSM7695928 | MV411_ATACseq_1h_DMSO_rep2 | 15019014 | 92.57 | 25222584 | 17419360 | 0.31 | 13.62 |
| GSM7695929 | MV411_ATACseq_1h_DMSO_rep3 | 15889335 | 92.84 | 26847784 | 18569499 | 0.31 | 13.49 |

#### Table S4. Mapping Metrics for mESC BRG1 Inhibitor Treated MNase-seq Samples from (Iurlaro, et al., 2021)

| Samples | SampleInformation | TotalReads | MappingRatio(%) | totalMappedPETs (MAPQ>=10) | uniquePETs | redundancy | fragmentLengthMean | fragmentLengthStd | finalUniqueReadsRemoveBlacklist | yield |
|---|---|---|---|---|---|---|---|---|---|---|
| GSM4798115 | MNase_DMSO_1 | 72539433 | 96.65 | 61516526 | 58481089 | 0.05 | 140.27 | 33.90 | 57106816 | 0.79 |
| GSM4798116 | MNase_DMSO_2 | 74014054 | 96.2 | 62288156 | 59168806 | 0.05 | 141.59 | 36.70 | 57758467 | 0.78 |
| GSM4798117 | MNase_BRM014-10uM_1 | 57578034 | 98.45 | 50623669 | 48405771 | 0.04 | 129.60 | 23.94 | 47342861 | 0.82 |
| GSM4798118 | MNase_BRM014-10uM_2 | 71671377 | 97.1 | 61274946 | 58259451 | 0.05 | 134.78 | 30.46 | 56910598 | 0.79 |

#### Table S5. Mapping Metrics for Published ChIP-seq Samples from (Egan, et al., 2016) and (Patel, et al., 2024)

| map to fly (spike-in) | SampleInformation | TotalReads | Mapping Ratio(%) | totalMappedReads | uniqueReads | redundancy | tssEnrichmentScore |
|---|---|---|---|---|---|---|---|
| GSM1890164 | PC9_EZH2inh_H3K4me3_Dmspike | 26911224 | 26.17 | 6060582 | 1864308 | 0.69 | 3.09 |
| GSM1890165 | PC9_control_H3K27me3_Dmspike | 42232080 | 7.33 | 2200997 | 2071306 | 0.06 | 1.51 |
| GSM1890166 | PC9_EZH2inh_H3K27me3_Dmspike | 37704531 | 27.56 | 7510098 | 6855981 | 0.09 | 1.29 |
| GSM1890167 | PC9_control_H3K4me3_Dmspike | 20602547 | 23.98 | 4252353 | 1219477 | 0.71 | 2.97 |
| map to human | | | | | | | |
| GSM1890164 | PC9_EZH2inh_H3K4me3_Dmspike | 26911224 | 71.51 | 16691327 | 5495172 | 0.67 | 23.82 |
| GSM1890165 | PC9_control_H3K27me3_Dmspike | 42232080 | 92.98 | 32600655 | 31236761 | 0.04 | 1.34 |
| GSM1890166 | PC9_EZH2inh_H3K27me3_Dmspike | 37704531 | 72.55 | 22117562 | 20429057 | 0.08 | 1.51 |
| GSM1890167 | PC9_control_H3K4me3_Dmspike | 20602547 | 73.03 | 13079959 | 4099482 | 0.69 | 21.02 |
| map to yeast (spike-in) | | | | | | | |
| GSM8439484 | HelaS3, mitotic, DMSO, 0.00025x yeast spike-in, H3K9ac | 8343480 | 2.68 | 169541 | 162405 | 0.04 | 1.09 |
| GSM8439486 | HelaS3, mitotic, DMSO, 0.0025x yeast spike-in, H3K9ac | 9960556 | 1.28 | 81953 | 76425 | 0.07 | 1.18 |
| GSM8439488 | HelaS3, mitotic, DMSO, 0.025x yeast spike-in, H3K9ac | 11643652 | 5.4 | 538061 | 519590 | 0.03 | 1.13 |
| GSM8439490 | HelaS3, mitotic, DMSO, 0.25x yeast spike-in, H3K9ac | 12300781 | 24.28 | 2703046 | 2463122 | 0.09 | 1.11 |
| GSM8439492 | HelaS3, mitotic, DMSO, 2.5x yeast spike-in, H3K9ac | 19567820 | 30.16 | 5339709 | 4577292 | 0.14 | 1.09 |

| | | | | | | | |
|---|---|---|---|---|---|---|---|
| GSM8439494 | HelaS3, mitotic, TSA, 0.00025x yeast spike-in, H3K9ac | 10677723 | 1.87 | 135894 | 127216 | 0.06 | 1.14 |
| GSM8439496 | HelaS3, mitotic, TSA, 0.0025x yeast spike-in, H3K9ac | 17786080 | 1.14 | 109972 | 96464 | 0.12 | 1.22 |
| GSM8439498 | HelaS3, mitotic, TSA, 0.025x yeast spike-in, H3K9ac | 11430666 | 4.78 | 447830 | 431278 | 0.04 | 1.09 |
| GSM8439500 | HelaS3, mitotic, TSA, 0.25x yeast spike-in, H3K9ac | 20713555 | 14.21 | 2613789 | 2455841 | 0.06 | 1.11 |
| map to human | | | | | | | |
| GSM8439484 | HelaS3, mitotic, DMSO, 0.00025x yeast spike-in, H3K9ac | 8343480 | 97.08 | 7342268 | 7245589 | 0.01 | 6.32 |
| GSM8439486 | HelaS3, mitotic, DMSO, 0.0025x yeast spike-in, H3K9ac | 9960556 | 97.55 | 8788099 | 8644047 | 0.02 | 6.19 |
| GSM8439488 | HelaS3, mitotic, DMSO, 0.025x yeast spike-in, H3K9ac | 11643652 | 93.01 | 9720783 | 9561352 | 0.02 | 5.17 |
| GSM8439490 | HelaS3, mitotic, DMSO, 0.25x yeast spike-in, H3K9ac | 12300781 | 74.72 | 8292870 | 8156357 | 0.02 | 6.14 |
| GSM8439492 | HelaS3, mitotic, DMSO, 2.5x yeast spike-in, H3K9ac | 19567820 | 69.67 | 11970438 | 11728645 | 0.02 | 2.31 |
| GSM8439494 | HelaS3, mitotic, TSA, 0.00025x yeast spike-in, H3K9ac | 10677723 | 97.96 | 9461917 | 9324459 | 0.01 | 5.67 |
| GSM8439496 | HelaS3, mitotic, TSA, 0.0025x yeast spike-in, H3K9ac | 17786080 | 98.76 | 15854488 | 15605344 | 0.02 | 5.07 |
| GSM8439498 | HelaS3, mitotic, TSA, 0.025x yeast spike-in, H3K9ac | 11430666 | 95.16 | 9770364 | 9623468 | 0.02 | 5.27 |
| GSM8439500 | HelaS3, mitotic, TSA, 0.25x yeast spike-in, H3K9ac | 20713555 | 85.6 | 15936259 | 15685187 | 0.02 | 4.60 |

#### Table S6. Summary of Deposited Data and Used Public Data

| Samples | Source | GEO accessions |
|---|---|---|
| DNase-seq of mouse DN3 wild-type and GATA3 knockout cells | This manuscript | GSE300647 |
| DNase-seq of mouse fibroblast wild-type and BRG1-AID cells (with human 293T cells as spike-in) | This manuscript | GSE300647 |
| GATA3 ChIP-seq from mouse DN cells | (Wei, et al., 2011) | GSM523221 |
| BRG1 ChIC-seq from mouse fibroblast cells | (Ren, et al., 2024) | GSM7713390<br>GSM7713391 |
| DNase-seq of mouse embryonic stem cells (mESC) | (Yue, et al., 2014) | GSM1014154 |
| CUT&RUNG of mESC BRG1 dTAG cells | (Hagihara, et al., 2025) | GSM8447862<br>GSM8447863<br>GSM8447864<br>GSM8447865<br>GSM8447866<br>GSM8447867<br>GSM8447868<br>GSM8447869<br>GSM8447870<br>GSM8447871<br>GSM8447872<br>GSM8447873 |

| | | |
|---|---|---|
| ATAC-seq of mESC BRG1 dTAG cells | (Hagihara, et al., 2025) | GSM8447850<br>GSM8447851<br>GSM8447852<br>GSM8447853<br>GSM8447854<br>GSM8447855<br>GSM8447856<br>GSM8447857<br>GSM8447858<br>GSM8447859<br>GSM8447860<br>GSM8447861 |
| ATAC-seq of human MV411 cells treated with DMSO, AU15330, and BRM014 (1-hour treatment with Drosophila melanogaster S2 cells as spike-in) | (Martin, et al., 2023) | GSM7695921<br>GSM7695922<br>GSM7695923<br>GSM7695924<br>GSM7695925<br>GSM7695926<br>GSM7695927<br>GSM7695928<br>GSM7695929 |
| MNase-seq of mouse embryonic stem cells (mESC) treated with DMSO and BRM014 | (Iurlaro, et al., 2021) | GSM4798115<br>GSM4798116<br>GSM4798117<br>GSM4798118 |
| DNase-seq of mouse embryonic stem cells (mESC) | (Yue, et al., 2014) | GSM1014154 |
| H3K4me3 and H3K27me3 ChIP-seq of PC9 cells treated with DMSO and EZH2 inhibitor | (Egan, et al., 2016) | GSM1890164<br>GSM1890165<br>GSM1890166<br>GSM1890167 |
| H3K9ac ChIP-seq of HeLa-S3 cells with variable ratios of yeast spike cells | (Patel, et al., 2024) | GSM8439483<br>GSM8439484<br>GSM8439485<br>GSM8439486<br>GSM8439487<br>GSM8439488<br>GSM8439489<br>GSM8439490<br>GSM8439493<br>GSM8439494<br>GSM8439495<br>GSM8439496<br>GSM8439497<br>GSM8439498<br>GSM8439499<br>GSM8439500 |
| ATAC-seq and RNA-seq of CD8+ cells comparing resting vs. stimulated conditions | (Shan, et al., 2022) | GSM5373919<br>GSM5373920<br>GSM5373923<br>GSM5373924<br>GSM5373928<br>GSM5373929<br>GSM5373933<br>GSM5373934 |

### Figure S1

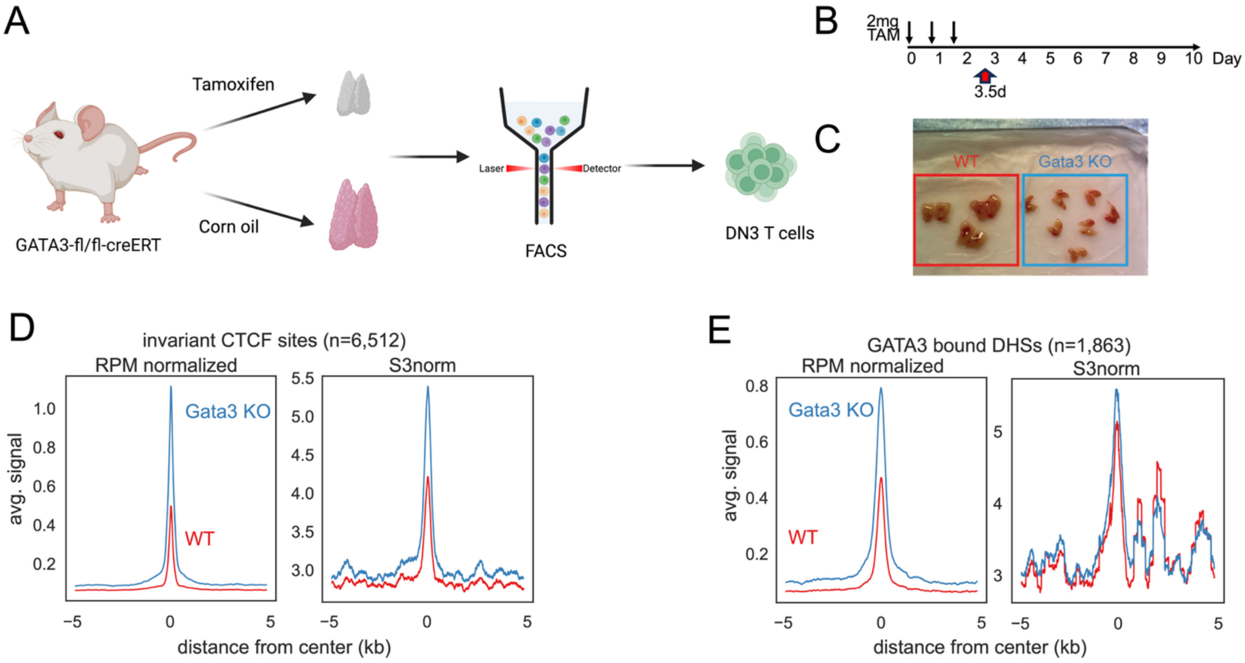

F

| | Run time | Max memory | input | output |
|---|---|---|---|---|
| Ryder | 328.61 s | ~1.23 GiB | RPM normalized bigWig files | normalized bigWig file |
| S3norm | 767.38 s | ~6.15 GiB | counts in bin as bedGraph files | normalized bedGraph file |

Note: Ryder is parallel designed and only run with 1 CPU comparing with S3norm

**Figure S1.** Experimental design for Gata3 knockout DN3 DNase-seq and comparison with S3norm normalization.

(A) Schematic of the experimental strategy for generating Gata3 knockout DN3 thymocytes. Gata3 fl/fl -CreERT mice were treated with tamoxifen to induce Gata3 deletion or with corn oil as control. DN3 thymocytes were isolated by FACS for DNase-seq analysis.

(B) Tamoxifen treatment schedule. Mice received 2 mg tamoxifen for three consecutive days, and DN3 cells were collected 3.5 days after the first injection.

(C) Representative thymus images from control and Gata3 knockout mice after treatment, illustrating the developmental effect of Gata3 deletion.

(D) Aggregate DNase-seq profiles centered on invariant CTCF-associated DHSs in WT and Gata3 KO DN3 cells. Left, RPM-normalized signals; right, S3norm-normalized signals. Invariant CTCF-associated DHSs were used as stable reference regions to evaluate normalization performance.

(E) Aggregate DNase-seq profiles centered on GATA3-bound DHSs in WT and Gata3 KO DN3 cells. Left, RPM-normalized signals; right, S3norm-normalized signals.

(F) Runtime and memory comparison between Ryder and S3norm for the Gata3 WT versus KO DNase-seq normalization benchmark. Ryder was run using one CPU for comparison with S3norm. Input and output formats for each method are indicated. Ryder used RPM-normalized BigWig files as input and generated normalized BigWig output, whereas S3norm used binned count bedGraph files as input and generated normalized bedGraph output.

### Figure S2

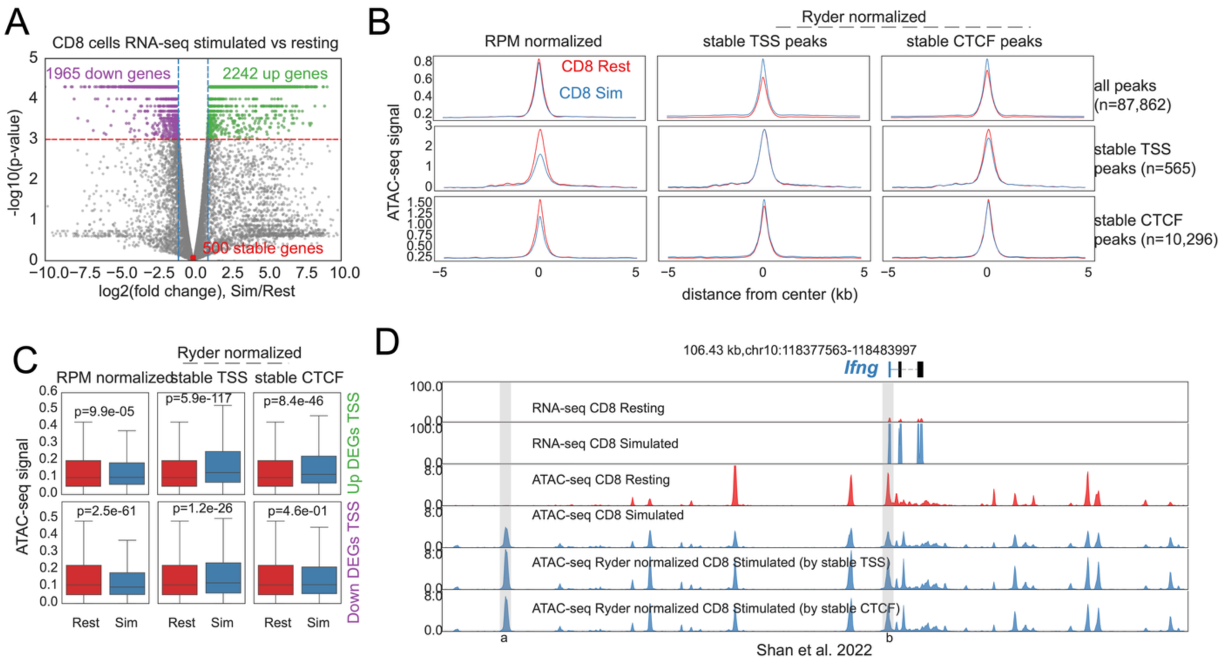

**Figure S2. Stable TSSs can serve as alternative internal reference regions for Ryder normalization in CD8 T cell activation.**

(A) Volcano plot of RNA-seq differential gene expression between stimulated and resting CD8 T cells. Upregulated genes, downregulated genes, and stable genes are indicated. A set of 500 stable genes was selected, as the genes in both condition RPKM > 1 and least absolute fold change, to define stable transcription start sites (TSSs) for alternative Ryder internal-reference normalization. Data were reanalyzed from (Shan, et al., 2022). Differentially expressed genes, fold change and P-values were obtained by Cuffdiff (Trapnell, et al., 2013).

(B) Aggregate ATAC-seq profiles in resting and stimulated CD8 T cells centered on all ATAC-seq peaks, stable TSSs, and stable CTCF-associated peaks. Left column, RPM-normalized ATAC-seq signals; middle column, Ryder-normalized signals using stable TSSs as internal reference regions; right column, Ryder-normalized signals using stable CTCF-associated peaks as internal reference regions. Data were reanalyzed from (Shan, et al., 2022). Stable TSSs are defined as the ATAC-seq peaks overlapped with the 500 stable expressed genes' TSS. Stable CTCF are defined as the overlapped peaks between resting and stimulated CD8+ cells, and further overlapped with in-variant CTCF sites we curated.

(C) Boxplots comparing ATAC-seq signals at TSSs of upregulated and downregulated genes in resting and stimulated CD8 T cells. Signals are shown for RPM normalization, Ryder normalization using stable TSSs, and Ryder normalization using stable CTCF-associated peaks. P-values above comparisons were calculated across matched TSS regions using a two-sided paired Wilcoxon signed-rank test.

(D) Genome browser view of the *Ifng* locus showing RNA-seq and ATAC-seq signals in resting and stimulated CD8 T cells. Ryder-normalized ATAC-seq tracks using either stable TSSs or stable CTCF-associated peaks preserve stimulation-induced accessibility increases at the *Ifng* locus.

### Figure S3

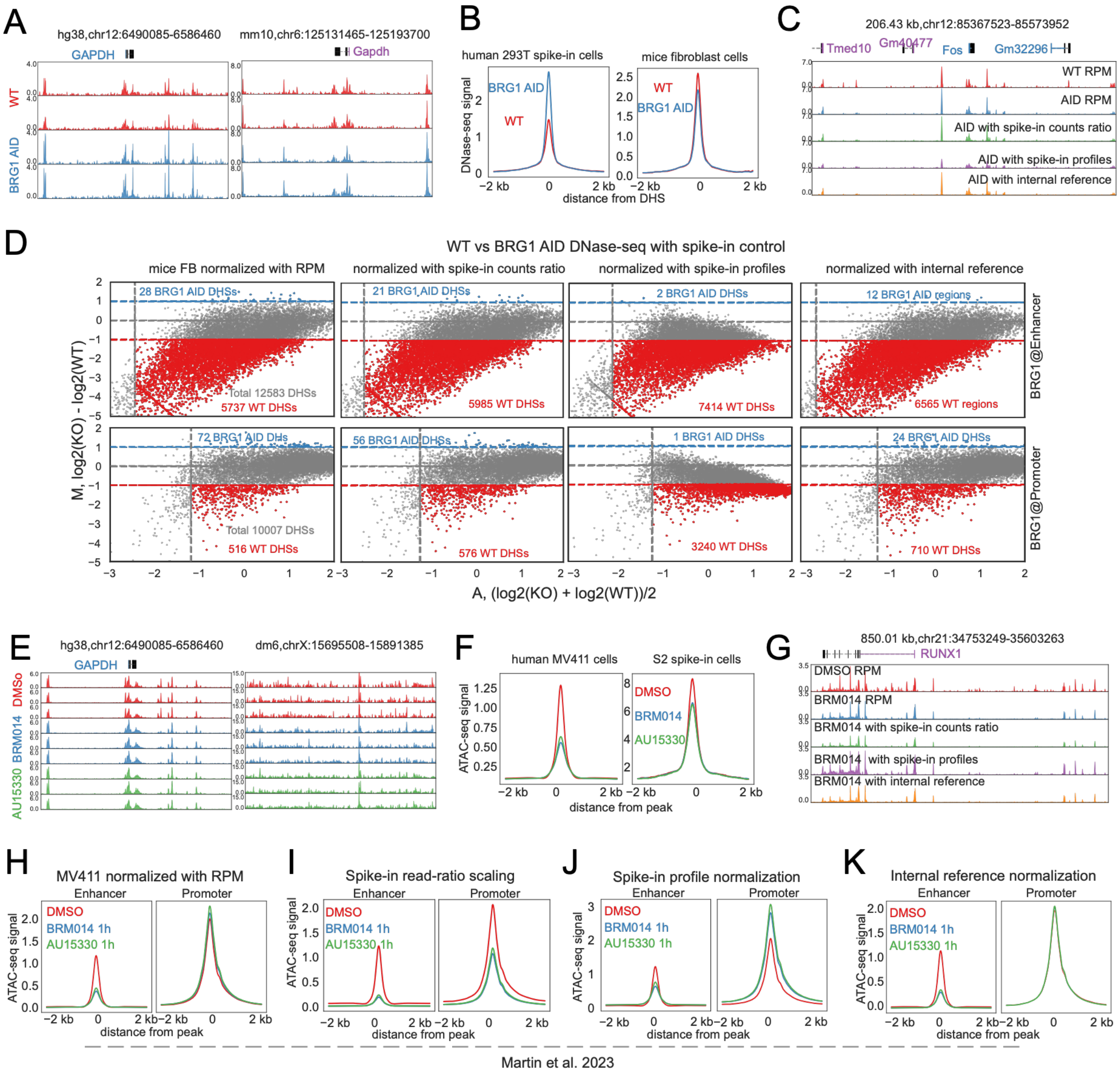

**Figure S3. Spike-in and internal-reference normalization comparisons in BRG1 perturbation DNase-seq and ATAC-seq datasets.**

(A) Genome browser views of representative gene loci showing DNase-seq signals (reads per million) in WT and BRG1-AID samples before normalization. Presented data were generated in this study.

(B) Aggregate plots of DNase-seq signals centered on all identified DNase Hypersensitive Sites (DHSs) in mouse (left) and human spike-in (right) cells.

(C) Genome browser view comparing the effect of different normalization methods on DNase-seq signal in WT versus BRG1-AID cells. Methods shown are standard RPM, spike-in read-count ratio, Ryder using the spike-in profile, and Ryder using internal reference regions.

(D) MA-plots showing differential BRG1-bound DHSs (enhancers and promoters) between WT and BRG1-AID cells. The comparison illustrates how the number and distribution of significant sites change across different normalization strategies.

(E) Genome browser tracks of ATAC-seq data from human MV411 cells treated with DMSO, BRM/BRG1 inhibitor AU15330, or BRM014, using Drosophila S2 cells as spike-in controls (Data from (Martin, et al., 2023)).

(F) Aggregated ATAC-seq signals at identified peaks in Drosophila and human cells prior to normalization.

(G) Genome browser views of ATAC-seq data comparing MV411 cells treated with DMSO or BRM014 under various normalization strategies.

(H) Aggregate ATAC-seq signals on human MV411 cells ATAC-seq peaks, comparing treatment of DMSO, AU15330 and BRM014, which are BRG1 and BRM inhibitors, separated into enhancers and promoters. Signals normalized with RPM. Data from (Martin, et al., 2023).

(I) Same as (H), but signals further scaled by the ratio of spike-in read counts between samples.

(J) Same as (J), with additional normalization using spike-in signal profiles: S2 ATAC-seq peaks normalized to derive parameters applied to mouse data.

(K) Same as (J), with further normalization using internal reference regions—MV411 ATAC-seq peaks overlapping human invariant CTCF sites (Fang, et al., 2020).

#### Figure S4

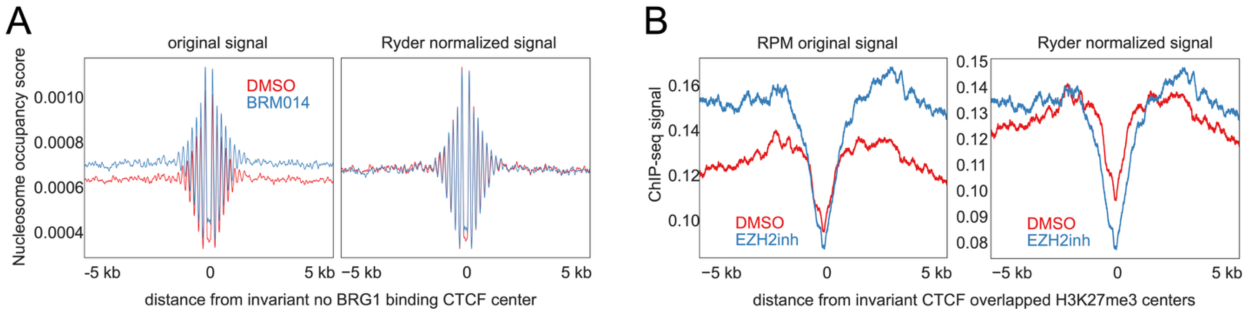

**Figure S4. Ryder normalization aligns internal reference regions in MNase-seq and H3K27me3 ChIP-seq datasets.**

(A) Aggregate plots of MNase-seq signal centered on invariant CTCF sites in mESCs treated with DMSO or a BRG1 inhibitor (BRM014). The comparison shows raw RPM signals (top) versus Ryder-normalized signals (bottom).

(B) Aggregate plots of H3K27me3 ChIP-seq signal centered on invariant CTCF sites in PC9 cells treated with DMSO or an EZH2 inhibitor (GSK126). The comparison highlights the difference between raw RPM signals (top) and Ryder-normalized signals (bottom) in a global-decrease scenario.
